# Dual RNAseq of human leprosy lesions identifies bacterial determinants linked to host immune response

**DOI:** 10.1101/354407

**Authors:** Dennis Montoya, Priscila R. Andrade, Bruno J. A. Silva, Rosane M. B. Teles, Bryan Bryson, Saheli Sadanand, Teia Noel, Jing Lu, Euzenir Sarno, Kristine B. Arnvig, Douglas Young, Ramanuj Lahiri, Diana L. Williams, Sarah Fortune, Barry R. Bloom, Matteo Pellegrini, Robert L. Modlin

## Abstract

To understand how the interaction between an intracellular bacterium and the host immune system contributes to outcome at the site of infection, we studied leprosy, a disease that forms a clinical spectrum, in which progressive infection by the intracellular bacterium *Mycobacterium leprae* is characterized by the production of type | IFNs and antibody production. We performed dual RNAseq on patient lesions, identifying a continuum of distinct bacterial states that are linked to the host immune response. The bacterial burden, represented by the fraction of bacterial transcripts, correlates with a host type | IFN gene signature, known to inhibit antimicrobial responses. Second, the bacterial transcriptional activity, defined by the bacterial mRNA/rRNA ratio, links bacterial heat shock proteins with the BAFF-BCMA host antibody response pathway. Our findings provide a platform for interrogation of host and pathogen transcriptomes at the site of infection, allowing insight into mechanisms of inflammation in human disease.

## Introduction

The interactions between the host immune response and an invading pathogen dictates the pathogenesis of an infectious disease. The spectrum of such interactions can be investigated in leprosy, caused by the intracellular bacterium *Mycobacterium leprae*, in which the clinical manifestations correlate with the host immune response to the pathogen (Ridley and Jopling, 1966). In the lesions of the self-limiting tuberculoid form (T-lep), bacilli are rare, and the immune response is characterized by a CD4^+^ T cell infiltrate (Modlin et al., 1982) and a IFN-γ transcriptional signature resulting in an effective antimicrobial responses in macrophages (Fabri et al., 2011; Montoya et al., 2009; Yamamura et al., 1991). By contrast, the lesions of the disseminated lepromatous form (L-lep) are characterized by abundant bacilli, B cells or plasma cells (Iyer et al., 2007; Ochoa et al., 2010), and expression of a type | interferon (IFN) gene program that suppresses macrophage antimicrobial activity (Teles et al., 2013). Some patients undergo a reversal reaction (RR) which manifests clinically as an upgrade from the L-lep to the T-lep form of the disease, associated with a change from a type | to type II IFN response.

Although *M. leprae* was the first human pathogen identified under the microscope, it has not been possible to grow this bacterium in vitro nor to develop a genetic system to study specific genes. Furthermore, research has been limited by the absence of animal models which mimic the clinical and immunologic spectrum of human disease (Kirchheimer and Storrs, 1971; Lahiri et al., 2005; Madigan et al., 2017a; Madigan et al., 2017b). Therefore, the study of human skin lesions from leprosy patients provides an opportunity to investigate the *M. leprae* determinants of the host immune response at the site of infection.

The advent of a “dual-RNAseq” approach has recently been applied to simultaneously sequence the transcriptomes of both the host and pathogen organisms in vitro, with subsequent isolation of each transcriptome *in silico* through alignment of the respective genomes of the organisms (Damron et al., 2016; Niemiec et al., 2017; Nuss et al., 2017; Thanert et al., 2017; Wesolowska-Andersen et al., 2017; Westermann et al., 2016; Zimmermann et al., 2017). The main challenge to this approach had been the size difference between bacterial and mammalian transcriptomes, which can lead to poor coverage of bacterial pathogens. This challenge has been addressed by molecular enrichment of bacterial RNA or by investigating cell cultures and animal models in which the multiplicity of infection is high (Damron et al., 2016; Niemiec et al., 2017; Nuss et al., 2017; Thanert et al., 2017; Wesolowska-Andersen et al., 2017; Westermann et al., 2016; Zimmermann et al., 2017). However, dual-RNA-seq has yet to be carried out in lesions of a human disease. We hypothesized that the high bacterial burden in L-lep lesions would provide an ideal opportunity to employ a dual-RNA sequencing approach to investigate the host-pathogen interaction at the site of human infectious disease.

## Results

### In situ dual RNA sequencing of host-pathogen transcriptomes identifies a link between pathogen abundance and type | interferon signaling

To investigate the host-pathogen interaction at the site of mycobacterial infection, RNA sequencing was performed on the total RNA from 24 leprosy skin biopsy specimens (9 L-lep, 6 T-lep, 9 RR) (Figure 1A). Microbeads with consensus sequences to human and bacterial ribosomal RNA (rRNA) were utilized for depletion. The enriched mRNA was then converted into sequencing libraries by random hexamer priming, sequenced via standard Illumina protocol, and mapped to both the human and *M. leprae* genomes (see Methods). Hierarchical clustering of both human and *M. leprae* transcriptomes revealed a distinct gene expression profile in L-lep lesions compared to T-lep and RR lesions (Figure 1B). The co-clustering of T-lep and RR samples (which was also seen with human gene clustering alone) is consistent with their similar histologic features, as observed previously in microarray studies (Bleharski et al., 2003; Teles et al., 2013).

**Figure 1.**
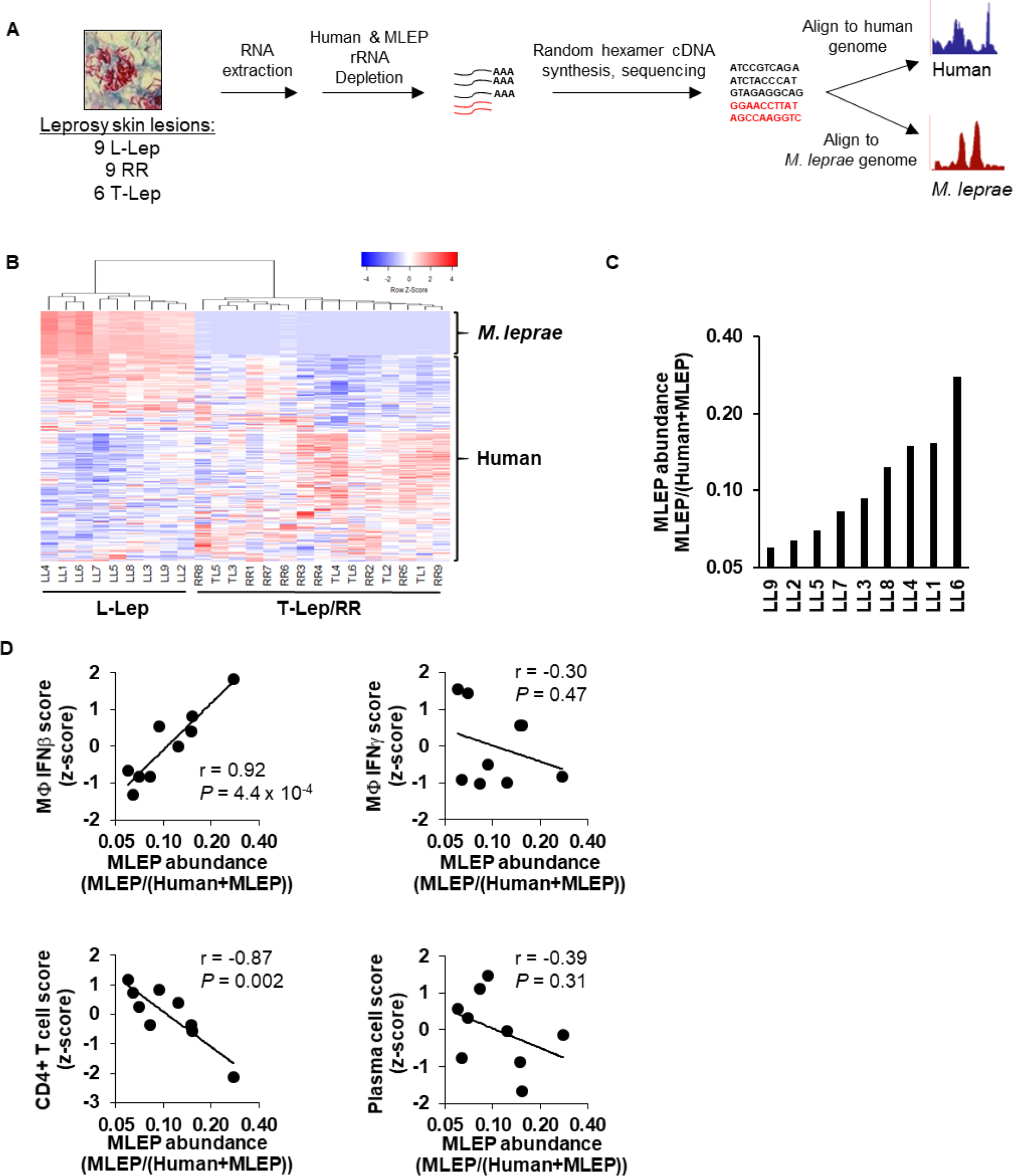
In situ dual-RNA sequencing of host-pathogentranscriptomes. **(A)** Experimental design fordual-RNAseq RNA extraction, library preparation, and sequence processing. **(B)** Unsupervised clustering analysis of both *M. leprae* and human gene expression across leprosy skin lesions (coefficient of variation > 1.0). Heatmap represents expression of genes (rows) and patients (columns) in which red indicates a higher and green a lower expression z-scoreper gene across patients. **(C)** Ratio of the sum of all *M. leprae* gene abundances over the total sum of human or *M. leprae* gene abundance across L-lep patient lesions. **(D)** Correlation plots of *M. leprae* abundance (log2-scale) against indicated SaVanT signature values (z-score)forlFN-β, IFN-γ, CD4+ T cells, or plasma cells per L-lep patient. *P*-value by student s t-test. See also Table S1 and Figure S1.

With a sequencing depth of ~50 million exonic reads, *M. leprae* reads were abundantly detected in the L-lep lesions (0.53-3.2 million exonic reads, Table S1) but at low levels in the RR or T-lep lesions (53-36,971 exonic reads), consistent with the histologic detection of bacilli in the different disease states (Ridley and Jopling, 1966). Reads mapping to the *M. leprae* genome had an average exome coverage of 22X and covered 95% of the total *M. leprae* genes (2618 genes) with at least five average exonic reads across L-lep samples. Furthermore, the fraction of the transcript abundance (which accounts for gene length, see Methods) mapping to *M. leprae* relative to abundance of transcripts mapping to either human or *M. leprae*, was variable across L-lep lesions and ranged from 0.06 to 0.38 (Figure 1C and Table S1). We next asked whether the variation in *M. leprae* abundance in the L-lep patients was linked to host immunologic features of the disease, through use of a gene signature-based analysis measuring the IFN-activation or cell type profile, which was used to deconvolute the cellular composition of each lesion (Lopez et al., 2017). We found a significant correlation of the *M. leprae* abundance with an IFN-β-activation gene expression signature (*r* = 0.92, *P* = 4.4 × 10^−4^), but not an IFN-γ signature (Figure 1D). By contrast, *M. leprae* abundance was not significantly correlated with a macrophage gene expression signature (Figure S1), suggesting that the type | IFN response is related to the bacterial load independently of macrophage numbers. Finally, we also observed that the *M. leprae* abundance was inversely correlated with the CD4^+^ T cell signature (r = −0.87, *P* = 1.1 × 10^−3^), but did not correlate with a plasma cell signature.

### Composition of the M. leprae transcriptome

We next investigated whether more specific properties of the bacterial transcriptome might be associated with immune signatures. Structural noncoding RNA, including transfer messenger RNA, ribonuclease P, and 23S ribosomal RNA (rRNA) accounts for 53.8% of the *M. leprae* transcriptome, while mRNA accounts for 46.2% (Table S2). Abundant mRNAs included virulence proteins that compose the ESX1 secretion system (esxA, esxB), the ESX1 associated proteins (the apparently operonic transcripts, espA, espC, espD MLBr00411 and PE3), and stress protein transcripts (cspA, hsp18, groEL2, groES), as well as the master transcriptional regulator, whiB1. Interestingly, many of the most abundant *M. leprae* RNAs were encoded by pseudogenes (Table S2), even though pseudogenes are not translated into full proteins due to nonsense mutations.

Although we depleted bacterial rRNA using affinity beads, we observed that certain regions of 23S rRNA were retained at high coverage, presumably because the rRNA was fragmented, and certain regions were not effectively captured by the beads, (Figure S2A). Sequencing a separate set of samples sequentially before and after rRNA depletion showed one 261 bp region in particular that was consistently retained (Figure S2B). We postulated that we could use this retained rRNA fragment as a surrogate for total 23S abundance. To confirm that the relative expression of this region matched 23S expression before rRNA depletion, cDNA, which was acquired from the nine L-lep lesions before depletion, was analyzed by qPCR. There was a high correlation (r = 0.71, *P* = 0.032) between the abundance of RNAseq 23S and that of the 261bp region (Figure S2C). As a control, *M. leprae* esxA an mRNA presumed to be unaffected by rRNA depletion, also showed significant correlation (r = 0.71, *P* = 0.014) before and after depletion.

### M. leprae transcriptional activity as a bacterial metric linked to a plasma cell response

The viability of *M. leprae* can be estimated from the ratio of rRNA transcripts to genomic DNA, an approach that leverages the difference in stability between rRNA and gDNA (Kralik et al., 2010; Liu et al., 2012; Martinez et al., 2009). Here we sought to extend this logic and estimate bacterial transcriptional activity based on measurements of the RNA only, exploiting the difference in mRNA and rRNA stabilities. For example, *in vitro*, mycobacterial mRNA has an average half-life of 9.5 minutes, while rRNA has an average half-life over 24h (Cangelosi and Brabant, 1997; Rustad et al., 2013). To validate that the mRNA:23S ratio correlates with the *M. leprae* transcriptional activity, we measured the ratio in *M. leprae* gene expression data from various conditions in which transcriptional activity is known to vary. *M. leprae* grown in the mouse footpad of immunocompromised (athymic nu/nu) mice were purified and placed into axenic culture for 0-96h. In these conditions *M. leprae* is unable to reproduce and its metabolism has been shown to slowly decrease (Davis et al., 2013). We calculated the transcriptional activity by dividing of the sum of the transcript abundance of all bacterial mRNA genes by the transcription abundance of 23S (based on the 1343455-1343551 region). As expected, we find that in these conditions, transcriptional activity, as measured by the mRNA:23S ratio, significantly decreased with time (*n* = 2, *P* = 0.005 at 48h, *P* = 0.006 at 96h) (Figure 2A). A second series of analyses was performed using data from conditions in which *M. tuberculosis* H37Rv was depleted of all nutrients and grown only in PBS, or *M. tuberculosis* H37Rv during log or transcriptionally quiescent stationary phase (Arnvig et al., 2011; Cortes et al., 2013). The mRNA:23S transcriptional activity measure was significantly lower in bacteria after nutrient depletion (*P* = 0.027), or in bacteria harvested in stationary phase compared to exponential phase (*P* = 0.002) (Figures S3A and S3B).

**Figure 2.**
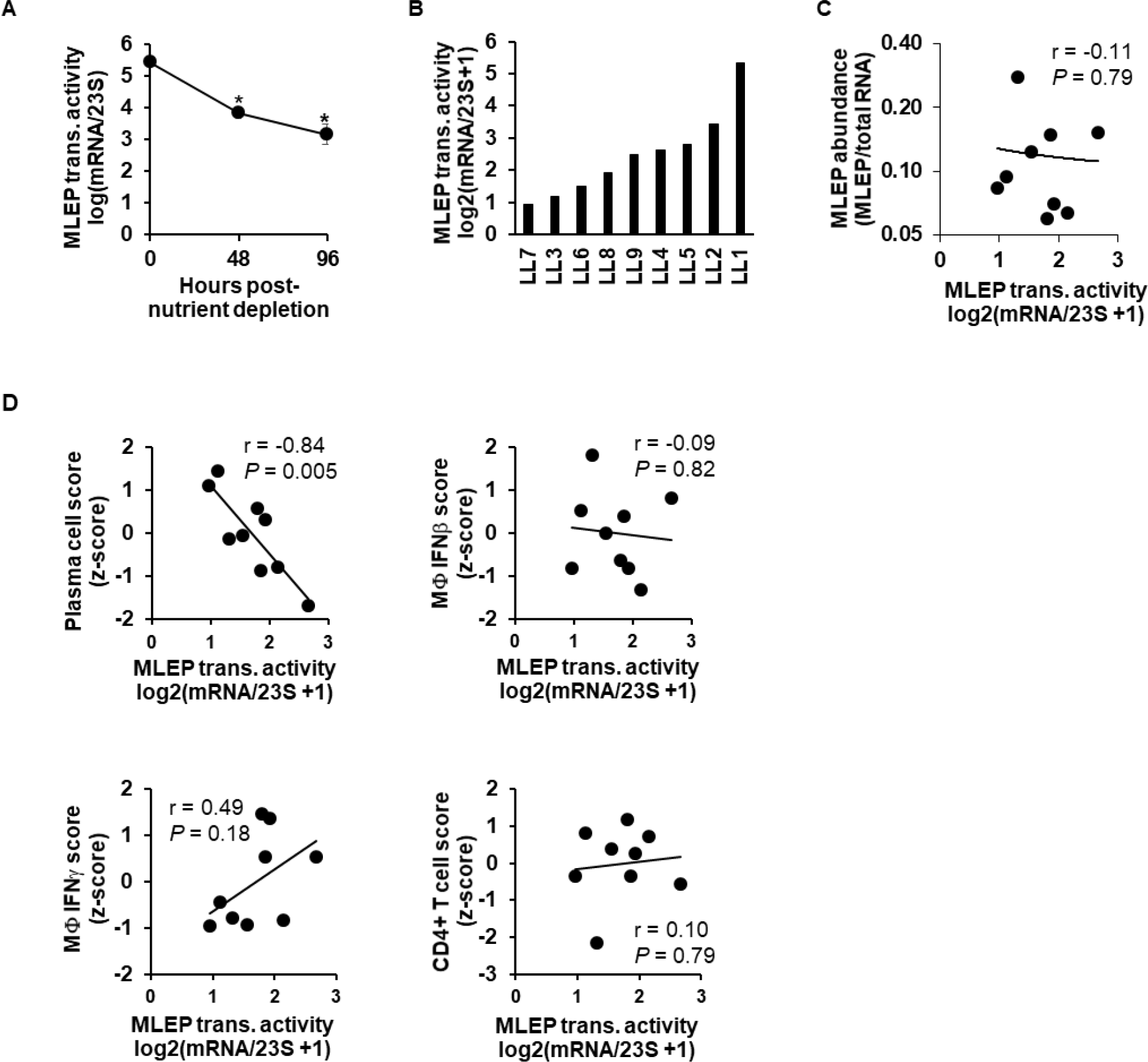
Measure of *M. leprae* transcriptional activity. **(A)** *M. leprae* derived from the footpad of *nu/nu* mice were isolated and cultured in vitro under starvation conditions for Oh, 48h, or 96h before RNA extraction and RNA sequencing. Data shown as mean transcriptional activity ± s.e.m. per timepoint (*n* = 2). \**P* < 0.05 by student’s t-test. **(B)** Ratio of the log2(x + 1) transformed sum of all *M. leprae* mRNA genes divided by 23S RNA abundance per L-lep lesion. **(C)** Plot of RNA abundance versus transcriptional activity *M. leprae* measures per L-lep patient. **(D)** Scatter plots of *M. leprae* transcriptional activity versus indicated SaVanT signature score per L-lep patient. *P*-value by student’s t-test. See also Table S2, Figure S2, and Figure S3.

Having established that the ratio of mRNA to rRNA provides a surrogate for ongoing bacterial transcriptional activity, we next asked if its variation across L-lep lesions correlated with host gene expression signatures that are characteristic of L-lep patients. This value was found to vary across L-lep lesions (Figure 2B and Table S1), and was largely independent of *M. leprae* abundance (Figure 2C, r = −0.11, *P* = 0.79). We found that the transcriptional activity of *M. leprae* was significantly negatively correlated with the plasma cell signature (*r* = -0.84, *P* = 0.005), but not the IFN-β, IFN-γ and CD4^+^ T cell signatures in L-lep lesions (Figure 2D).

### Identification of a M. leprae gene module associated with host antibody production in L-lep lesions

To identify the *M. leprae* gene modules associated with the variation in transcriptional activity, a weighted-gene correlation analysis (WGCNA) was performed on the bacterial transcriptome (Figure 3A) (Langfelder and Horvath, 2008b; Montoya et al., 2014). WGCNA is an unsupervised method that uses expression correlations to group genes into modules, similar to traditional clustering analysis, but also accounts for the network neighborhood of a given set of genes, thus lending more weight to gene networks with more reliable correlations. To account for the varying abundance of bacilli across patient lesions, the relative percentage of total bacterial mRNA was input for each *M. leprae* gene. We found that among the 14 modules that we identified, the *M. leprae* RNA abundance measure correlated most highly with the average transcriptional abundance (module eigengene) of the *MLEPpurple* module, although this did not reach significance (r = 0.5, *P* = 0.2). The *MLEPpurple* module contained two major components of the genes the ESX-1 secretion system, esxA and esxB, which are required for type | IFN induction in macrophages by mycobacterial infection (Stanley et al., 2007). The relative expression of esxA was significantly correlated with the type | IFN gene signature (r = 0.71, *P* = 0.03, Figure 3B). Meanwhile, the *M. leprae* transcriptional activity positively correlates with the *MLEPcyan* module eigengene, (r = 0.84, *P* = 0.005), and the *MLEPblue* module eigengene composed of 257 genes inversely correlates with transcriptional activity (r = −0.75, *P* = 0.02) (Figures 3A, 3C, and Table S3).

**Figure 3.**
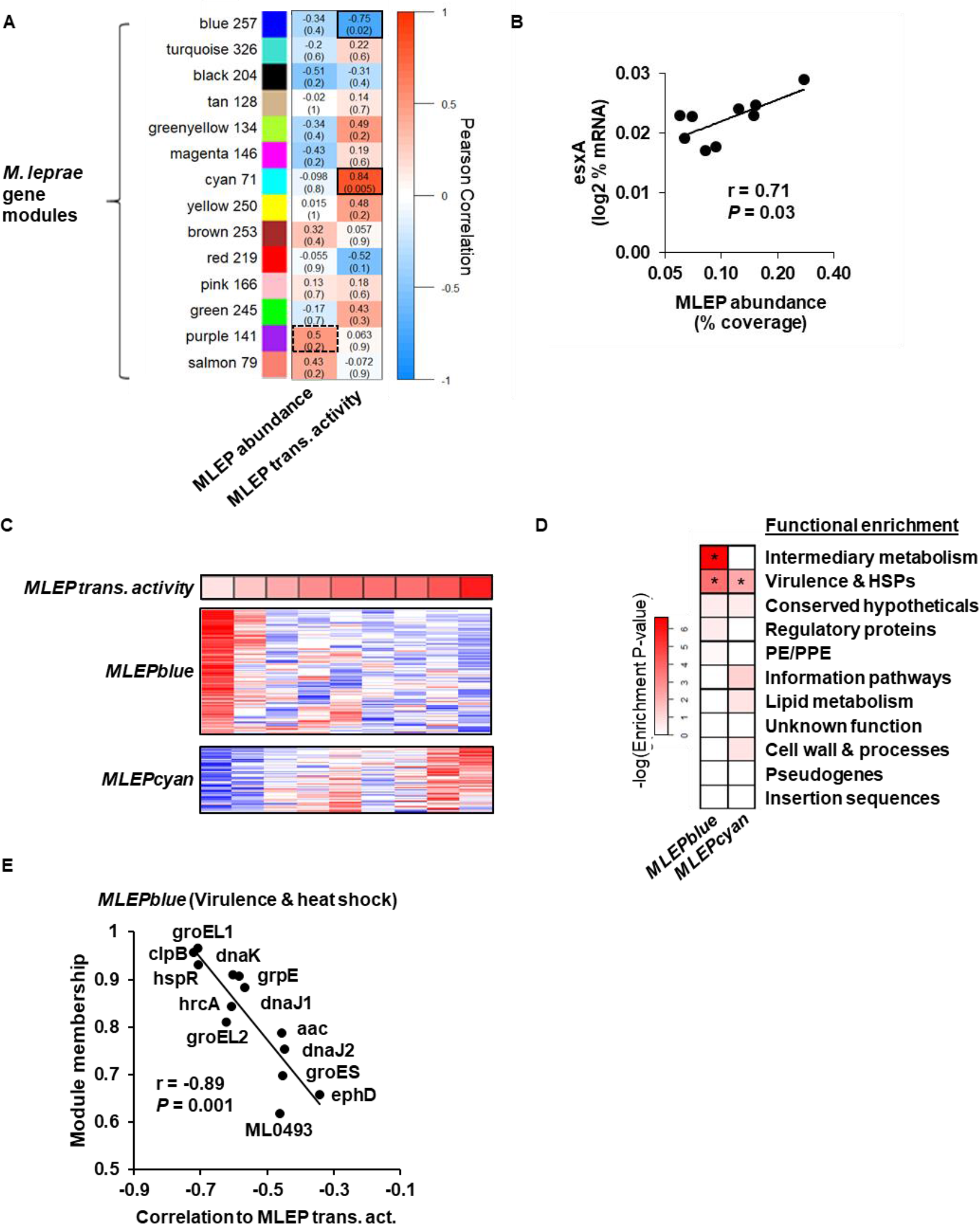
Module analysis of *M. leprae* transcriptome. **(A)** Unsupervised weighted gene-correlation analysis (WGCNA) of the *M. leprae* gene expression across L-lep patients. Heatmap represents the correlation of the module eigengene (rows) versus bacterial measure (columns) in which red indicates a positive and blue a negative Pearson correlation across patient samples. Number of genes per module indicated beside module color. Black outlines indicate highest and lowest correlation values. **(B)** Correlation plot of the *MLEPpurple* module gene, esxA, and the *M. leprae* abundance per L-lep patient. **(C)** Heatmap of *MLEPblue* or *MLEPcyan* module protein coding genes (pseudogenes excluded) across L-lep patients in order of *M. leprae* transcriptional activity (top row). High or low relative expression indicated by red or blue, respectively. **(D)** Functional category enrichment of genes within the *MLEPblue* or *MLEPcyan* modules. Heatmap color indicates the negative logarithm of the hypergeometric enrichment *P-*value per functional category. \**P* ≥ 0.05. **(E)** Plot of the WGCNA module membership and the correlation to the *M. leprae* transcriptional activity per gene from the *MLEPblue* ‘virulence and heat shock’ genes. *P*-value by student’s t-test. See also Table S3

The functional categories associated with the significantly correlated *M. leprae* gene modules were investigated using annotations from the Mycobrowser (Kapopoulou et al., 2011). The *MLEPcyan* and *MLEPblue* modules were significantly enriched in the ‘virulence, detoxification, adaptation’ functional category defined by Mycobrowser, but we noted there were numerous heat shock proteins (HSPs), hence we designate as ‘virulence and HSPs’ (Figure 3D). The *MLEPblue* module in particular was enriched in genes encoding HSPs, while *MLEPcyan* contained none. Furthermore, an additional enrichment in ‘intermediary metabolism’ was identified for the blue module. The correlation of individual *MLEPblue* virulence genes with *M. leprae* transcriptional activity correlated module membership, a measure of the genes most highly connected to the module (Figure 3E).

### Interactions between the M. leprae and host transcriptomes

To further understand the interaction between the bacterial and host transcriptomes in leprosy, the WGCNA module eigengenes derived from the *M. leprae* transcriptome and the metrics for bacterial abundance and transcriptional activity were correlated with the WGCNA modules derived from the human transcriptome within the same lesions. Of the 36 human modules generated, nine were highly correlated (*P* < 0.001) with the *M. leprae* transcriptional activity, abundance, or module eigengenes of the *MLEPblue* or *MLEPcyan* modules (Figure 4A). The most significant correlation was between the *HUMANgreenyellow* and the *MLEPblue* module eigengenes (r = 0.78, *P* = 7 × 10^−6^). Gene-Set Enrichment Analysis (GSEA) (Subramanian et al., 2007) of the *HUMANgreenyellow* module genes identified ‘humoral immune response’ as the top term, accounting for 27 of the 575 genes in the module. Overall, immunoglobulin genes accounted for 215 (37%) of the genes in module, including all of the of the immunoglobulin heavy chain constant genes (IGH) which determine antibody isotype (Figure 4B and Table S4). IGH transcripts were seven-fold more abundant in L-lep vs. T-lep lesions, in part due to the dominance of IGHM transcripts. Of the class switched isotypes, the total of the IGHG subtypes predominated in L-lep lesions followed in expression by IGHA (Figure 4B). Further investigation by Ingenuity Pathways Analysis (IPA) of *HUMANgreenyellow* indicated the canonical pathway most significantly enriched to be ‘Communication between Innate and Adaptive Immune Cells’ (hypergeometric enrichment *P* value = 1.71 × 10^−16^), identifying a pathway for immunoglobulin production. This pathway contains the ligands BAFF (B cell activating factor, aka TNFSF13B) and APRIL (a proliferation-inducing ligand, aka TNFSF13) which activates the B cell receptor BCMA (B cell maturation antigen, aka TNFRSF17), leading to production of IgG and IgA, all of which were differentially expressed in L-lep vs. T-lep lesions (Figure 4C). In addition to BCMA, the *HUMANgreenyellow* module genes IGHA1, IGHA2 and IGHG2 had significant inverse correlation with the *M. leprae* transcriptional activity (r = −0.84, *P* = 0.005; r = −0.79, *P* = 0.011; r = 0.76, *P* = 0.017, respectively, Figure 4D). BCMA inversely correlated with the *M. leprae* transcriptional activity (r = −0.77, *P* = 0.008), but positively correlated with the plasma cell signature (r = 0.96, *P* = 4 × 10^−5^) (Figures 4D, 4E, and Figure S4A).

**Figure 4.**
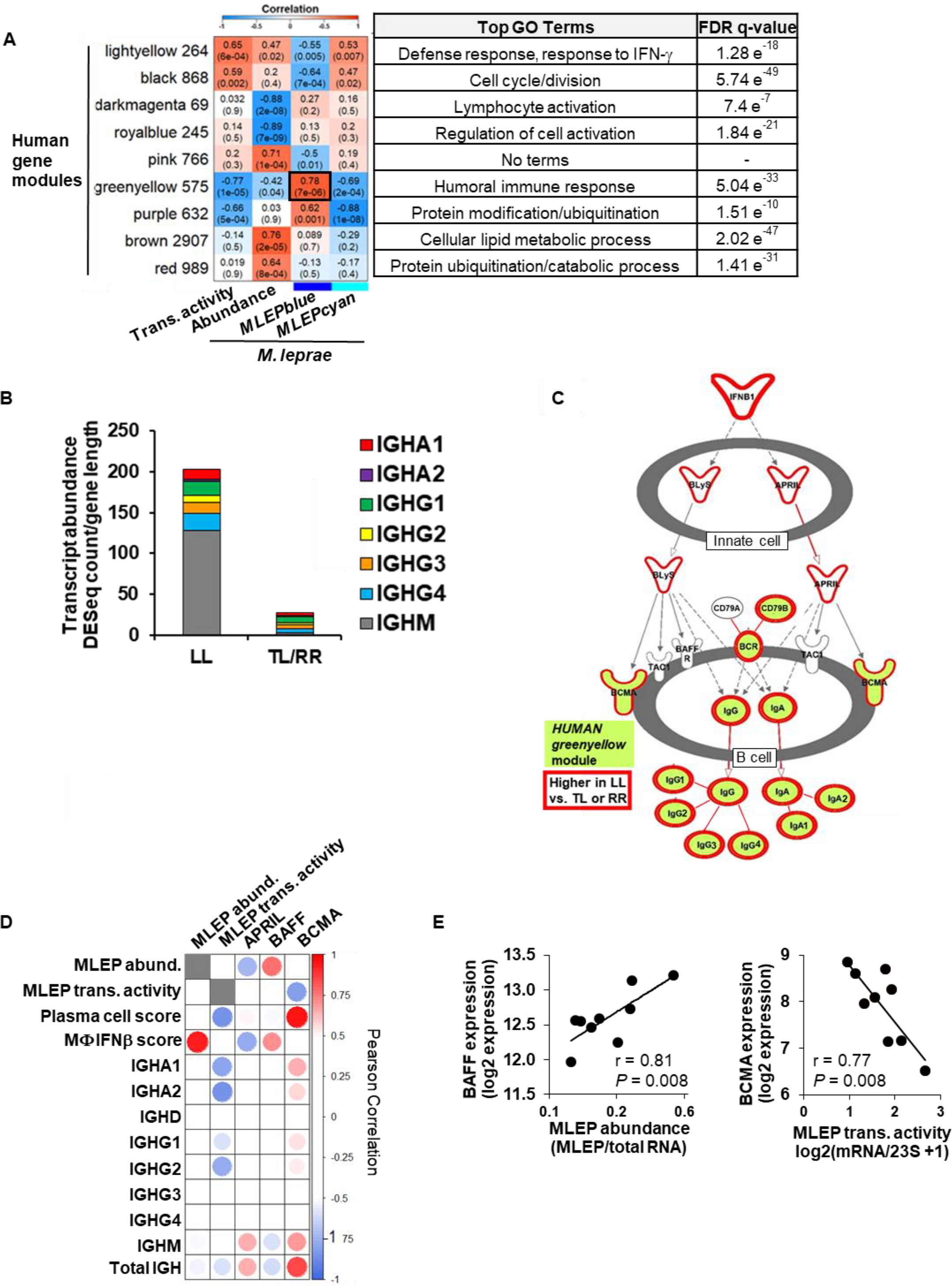
Interactions between host and pathogen transcriptomes. **(A)** WGCNA of the human gene expression across L-lep patients. Heatmap represents the correlation of the host module eigengene (rows) versus bacterial measure (columns) in which red indicates a positive and blue a negative Pearson correlation across patient samples. Only modules with a significant correlation (*P* < 0.001) in at least one bacterial measure or module eigengene are shown. Black outline indicates most significant positive correlation. Number of genes per module indicated beside module color and top gene ontology terms for module genes listed on right with the FDRq-value for enrichment per module. **(B)** Mean abundance of IGH immunoglobulin constant regions per leprosy clinical subtype. **(C)** Ingenuity Pathways canonical pathway ‘Communication between Innate and Adaptive immune cells’. Only section of pathway containing enriched genes shown. *HUMANgreenyellowmo6u\e* genes shaded green and genes significantly higher in LL vs TL/RR patients (fold change ≥ 1.5, P_adj_ ≤ 0.05 outlined in red. Bonferoni-adjusted *P* value by Wald test). **(D)** Correlation heatmap of BAFF and APRIL ligands and BCMA receptor expression versus *M. leprae* bacterial measures, human SaVanT plasma cell and IFN-β signature score, and individual immunoglobulin IGH constant region genes across L-lep patients. High or low Pearson correlation indicated by red or blue, respectively. **(E)** Correlation plots of *M. leprae* bacterial measures across L-lep skin lesions versus BAFF or BCMA expression. *P*-value by student’s t-test. See also Figure S4, Figure S5, and Table S4.

Although BAFF and APRIL were not present in the *HUMANgreenyellow*, they were significantly higher in L-lep versus T-lep or RR patients (1.7 fold-change, *P*_adj_ = 0.004, 1.6 fold-change, *P*_adj_ = 0.007, respectively). Although APRIL was expressed in L-lep lesions, it did not correlate with either *M. leprae* transcriptional activity or abundance. By contrast, BAFF expression significantly correlated with the *M. leprae* abundance (r = 0.81, *P* = 0.008), but not transcriptional activity, the plasma cell signature, nor IGH expression (Figure 4D). BAFF expression did correlate with the type | IFN activation signature score (r = 0.71, *P* =0.032) across L-lep patients (Figures 4D, 4E and Figure S4B). *M. leprae* abundance is likely linked to BAFF expression through type | IFN production, since in vitro *M. leprae* infection of monocyte-derived macrophages induces type | IFN (Teles et al., 2013) and BAFF (Figure S5A). Lastly, type | IFN stimulation induces BAFF in uninfected macrophages (Figure S5B).

Since BCMA expression appears to be linked to antibody expression, we investigated how the *MLEPblue* ‘virulence and heat shock protein’ genes were individually connected with BCMA expression. Six *M. leprae* genes (aac, ML0493, clpB, groEL1, groEL2, and hspR) significantly correlated with the expression of BCMA, the plasma cell signature, IGHA1, IGHA2, IGHG2, and the total amount of IGH (Figures 5A). GroEL1 and groEL2 also particularly demonstrated a strong correlation with the class-switched IGHA1 and IGHG2 antibody production (Figures 5A).

**Figure 5.**
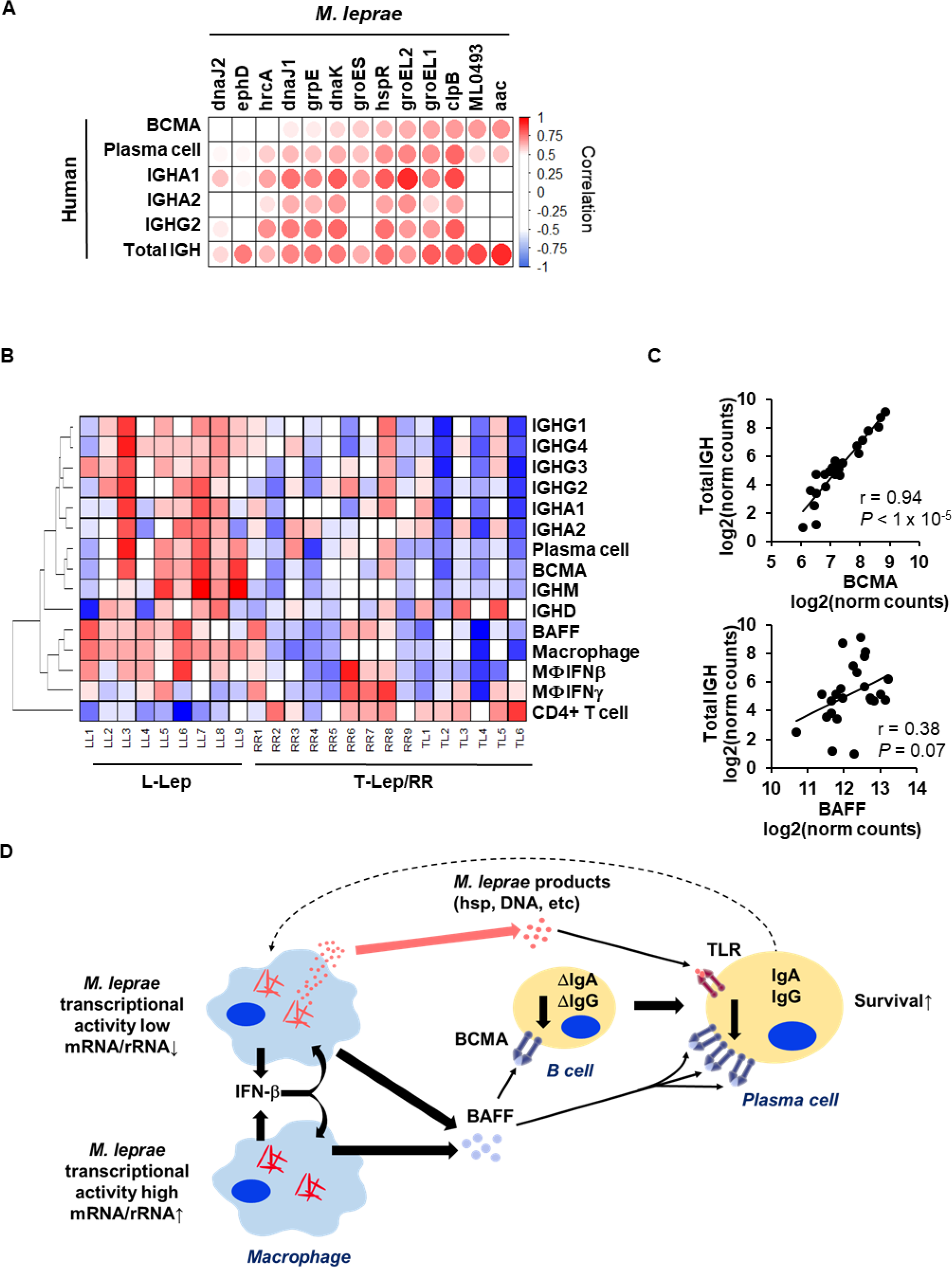
Bacterial stress proteins are linked to BAFF-BCMA induced antibody expression. **(A)** Correlation heatmap of *MLEPblue* ‘virulence and heat shock’ genes (columns) versus MLEP transcriptional activity and components of the host antibody response. Positive or negative Pearson correlation indicated by red or blue, respectively. **(B)** Heatmap of expression of human immune response components across the spectrum of leprosy clinical subtypes. Color represents the z-score across each row with red as high and blue as low relative expression. **(C)** Correlation plot of the total IGH abundance versus BCMA or BAFF expression across the spectrum leprosy samples (L-lep, T-lep, RR). *P*-value by student’s t-test. **(D)** Model for interaction between the pathogen and host humoral response. The model shows that bacterial abundance is correlated with induction oflFN-p leading to BAFF expression. At the same time, decreased *M. leprae* transcriptional activity is linked to the expression of *M. leprae* heat shock proteins, which can trigger TLR4 activation on plasma cells to upregulate the BAFF receptor, BCMA. The BAFF-BCMA interaction results in maturation and survival of class-switched antibody plasma cells.

To further understand the relevance of BAFF and BCMA to antibody responses in L-lep patients, we investigated the expression of key immune response genes and signatures across the leprosy disease spectrum. As previously stated, BAFF, BCMA, IGH as well as the individual immunoglobulin isoforms were more strongly expressed in L-lep vs. T-lep and RR lesions (Figure 5B). The plasma cell signature linked to BCMA and the type | IFN signature linked to BAFF were also greatest in L-lep lesions. Although BAFF and BCMA expression was highest in L-lep lesions, only the level of BCMA expression across all leprosy patients is tightly correlated with the total antibody expression (r = 0.94, P < 1 × 10^−6^, Figure 5B, C). Together, these data indicate that BCMA expression is central to immunoglobulin production at the site of disease in leprosy and is linked to decreased *M. leprae* transcriptional activity and a stress response gene module.

## Discussion

The locus of the battle between pathogens and host is primarily in tissue lesions, about which remarkably little is known for many pathogens. To define the molecular determinants of the host-pathogen interaction at the site of an easily-accessible mycobacterial infection, we performed dual RNAseq on leprosy lesions, simultaneously measuring both the human and *M. leprae* transcriptomes. The clinical spectrum of leprosy is defined by the number of skin lesions and microscopic quantification of bacilli, which grossly correlates with the host immunologic responses (Ridley and Jopling, 1966). This clinical spectrum defines a subgroup of patients, L-lep, presenting with disseminated lesions containing a high bacterial load. Here we define two independent molecular determinants of the pathogen state, each of which correlates with distinct aspects of the host immune response (Figure 5D). The first is the bacterial burden, represented by the fraction of bacterial transcripts, which correlates with a host type | IFN gene signature. We identified a second bacterial parameter, the transcriptional activity, defined by the *M. leprae* mRNA/rRNA ratio that is correlated with a plasma cell signature and antibody expression. Simultaneous measurement of both the human and *M. leprae* transcriptomes enabled us to identify associations between the host and pathogen, including a novel link between mycobacterial ‘stress response’ and ‘human antibody response’ gene networks, providing a precision medicine approach to identifying critical determinants of disease pathogenesis.

The current standard to assess leprosy bacterial burden in the clinical setting is the bacillary index (BI), developed 61 years ago, determined through enumeration of acid-fast stained bacteria per high power microscopic field from blood slit smears from either the earlobe or the disease lesion (Ridley, 1957). Here we developed a molecular definition of bacterial burden, measuring the *M. leprae* RNA abundance, finding that this metric correlated strongly with a host type | IFN signature. Previously, we found that IFN-β secretion by macrophages correlated with the multiplicity of infection by *M. leprae* in vitro (Teles et al., 2013). In addition, while the relative expression of bacterial esxA has been demonstrated in experimental models to be required for mycobacteria-induced type | IFN expression, we provide evidence linking *M. leprae* esxA to the human type | IFN signature at the site of infection. Although, the type | IFN inducible program is a major immune factor for control of viral infection, it has been shown to suppress antibacterial responses, particularly to intracellular infections (Boxx and Cheng, 2016; Teles et al., 2013).

Although humoral immune responses are robust in multibacillary L-lep patients, little is known about antibody production at the site of infection (Iyer et al., 2007; Ochoa et al., 2010). Here, we found that the plasma cell gene signature did not correlate with the *M. leprae* RNA abundance, but this measure does not distinguish active versus quiescent bacilli. We overcome this limitation by considering a second property of bacilli, their transcriptional activation state defined by the ratio of bacterial mRNA to rRNA. This ratio was found to be independent of the *M. leprae* RNA abundance. We find that the transcriptional activity of the bacilli in each individual is strongly correlated with the abundance of plasma cells, as measured by the plasma cell signature.

We found that WGCNA was a powerful approach to associate bacteria and host gene networks, uncovering a molecular mechanism that links *M. leprae* transcriptional activity to the plasma cell response. Low bacterial transcriptional activity correlated with an *M. leprae* network, *MLEPblue*, which was linked to a host gene module *HUMANgreenyellow* that was enriched for antibody genes and BCMA, a cell surface receptor on plasma cells known to induce IgA and IgG class-switched antibody production in B cells (Marsters et al., 2000). A ligand for BCMA, BAFF, correlated with *M. leprae* abundance and a type | IFN program. These data identify a mechanism connecting *M. leprae* abundance and transcriptional activity leading to the enhanced humoral response in L-lep patients. The regulation of BCMA expression is critical to the induction of this humoral response as the expression of BCMA, but not BAFF was tightly linked to the immunoglobulin expression across the spectrum of leprosy. Mining the correlated bacteria and human gene networks, we identified an association between gene expression of *M. leprae* heat shock proteins and BCMA. We suggest that *M. leprae* under immune attack are stressed and transcriptionally low, cutting back on protein synthetic functions and increasing the ratio of expression of proteostasis genes such as HSPs. Furthermore, some stressed bacteria are dying, releasing their major cytoplasmic and nuclear contents. This would include DNA (Kim et al., 2011), known to trigger TLR9 on B cells and plasma cells, leading to upregulation of BCMA thereby allowing the BCMA ligand BAFF to augment antibody production. Alternatively, the increased ratio of HSPs can be released by dying bacteria (Vargas-Romero et al., 2016) and subsequently released from dying macrophages or secreted in exosomes (Giri et al., 2010). Of these proteins, GroEL1/2 and DnaK, have been shown to activate TLR4 (Asea et al., 2002; Bulut et al., 2005), which is also present on plasma cells (Dorner et al., 2009). These heat shock proteins are also known to be major antigens for the antibody response (Young et al., 1988). We noted that GroEL2 expression had the strongest correlation with a IgA1 antibody response, characteristic of a mucosal immune response, perhaps reflecting nasopharyngeal involvement in leprosy patients (Barton, 1975).

Alternatively, the correlation between groEL1/2 expression and an antibody response in L-lep lesions may be part of a host defense response. In tuberculosis, antibodies, including IgA, have been shown to contribute to an antimicrobial response against mycobacteria (Lu et al., 2016; Zimmermann et al., 2016). Targeting of heat shock proteins has long been used for bacterial vaccine design and in particular, *M. leprae* GroEL2 (Lima et al., 2003; Lorenzi et al., 2010; Lowrie et al., 1999; Souza et al., 2008; Tascon et al., 1996); and intranasally-administered *M. tuberculosis* DnaK (Chuang et al., 2018) have been shown to engender protective cell-mediated and humoral immune responses to tuberculosis in mouse models. Our data demonstrate a correlation between decreased *M. leprae* transcriptional activity with bacterial heat shock protein expression and the host humoral response, suggesting further investigation of the efficacy of mycobacterial heat shock proteins in prophylactic vaccines. On the other hand, the link between *M. leprae* heat shock protein gene expression and the host antibody response may reflect pathogenesis of the infection, indicating release of bacterial products that contribute to immunopathology. Although the expression of BAFF, BCMA, and antibody genes varied in L-lep patients, the levels were generally higher than in T-lep or RR patients. Excessive BAFF signaling of B cells has been shown to increase autoantibody production (Gross et al., 2000; Mackay et al., 1999; Zhang et al., 2001), which are prominent in L-lep patients complicating the differential diagnosis of systemic lupus erythematosus and syphilis (de Larranaga et al., 2000; Loizou et al., 2003).

Characterization of the transcriptionally inactive bacterial program in vivo is relevant to antibiotic drug design as bacteria in stationary phase and/or metabolic quiescence can result in increased ‘persister’ subpopulations that are phenotypically tolerant but not genetically resistant to antibiotics (Eng et al., 1991; Kester and Fortune, 2014). In particular, mycobacterial groEL1 and dnaK have been shown to be actively transcribed in isoniazid-tolerant *M. tuberculosis* persisters and stationary phase. Mycobacteria transformed with a groEL promoter reporter construct have been shown to have very low levels of activity during the exponential phase and increase progressively during the late exponential and stationary phases. Current live/dead reporters of mycobacterial species use a groEL promoter to drive the Mcherry persistent response that represent dying or dead bacteria (Martin et al., 2012; Sille et al., 2011). Lastly, these proteins have been recently described as part of the mycobacterial proteostasis network composed of GroEL1/GroEL2, GroES, DnaK, ClpB, DnaJ1, DnaJ2, and GrpE (Lupoli et al., 2016) which can aid the bacterium in surviving protein damage due to antibiotic treatment or the host antibacterial response. However further study would be needed to determine if the transcriptional inactivity defined by a low bacterial mRNA:rRNA ratio is seen in these ‘persister’ and live/dead bacterial models.

Previous studies seeking to gain insight into the battle between the host and a pathogen through dual RNA-seq has been limited to the study of infected cells in tissue culture or in mouse models of infection, often using genetically modified bacteria or different mouse strains. It is difficult to translate this approach to human infectious disease given that both the pathogen and the host cannot be genetically modified. Here, for the first time, we were able to define host-pathogen transcriptional associations in lesions of a human disease. The complex cellular environment of tissue specimens provides a bioinformatics challenge to analysis. Previous dual RNAseq studies of in vitro infected cells have been limited to only one cell type while in vivo studies in mouse models lacked analysis of the tissue cellular environment (Damron et al., 2016; Nuss et al., 2017; Pittman et al., 2014; Thanert et al., 2017). Here, we developed a computational framework that reduces the each disease lesion into a series of scores representing the host cellular immune pathways and pathogen abundance or activity. This reduction enabled us to correlate the spectrum of interactions between the pathogen and host immune response, identifying human and bacterial gene modules that are co-regulated across patients. Together, the transcriptional state and bacterial burden that varied across patients provide separate but informative metrics about the pathogenesis of mycobacterial disease that are likely to be general for other intracellular pathogens, revealing that the immune system responds to bacterial states and not just their abundance. As such, this approach provides a paradigm for investigating additional host-pathogen interactions in human disease.

## Acknowledgements

Financial support from NIAID AAI15006-004 for provision of *M. leprae.* Funding provided by UCLA CTSI grant number UL1TR000124, NIAMS NIH3P50AR06302. This work used computational and storage services associated with the Hoffman2 Shared Cluster provided by UCLA Institute for Digital Research and Education’s Research Technology Group.

## Author Contributions

Conceptualization, D.J.M and R.L.M; Methodology, D.J.M and M.P.; Investigation: D.J.M, R.M.B.T, R.L., D.L.W., P.A.; Formal analysis: D.J.M., M.P., T.N. J.L.; Writing - Original Draft, D.J.M, R.L.M, M.P.; Writing - Review & Editing, D.J.M. R.L.M, M.P., S.S, B.B, K.B.A, D.Y., S.F., B.R.B.; Resources, E.S., K.B.A, R.L., D.L.W; Supervision: R.L.M., M.P.; Funding Acquisition: R.L.M, M.P., D.J.M.

## Declaration of Interests

None

## Materials and Methods

### Patients and clinical specimens

Patients with leprosy were classified according to the criteria of Ridley and Jopling (Ridley and Jopling, 1966); all T-lep patients classified as borderline tuberculoid, and all L-lep patients had lepromatous leprosy. All L-lep and T-lep skin biopsies were taken at the time of diagnosis before treatment, and reversal reaction biopsies were upgrading from patients originally diagnosed with borderline lepromatous leprosy, therefore starting from a different part of the disease spectrum than the L-lep group. Specimens were embedded in OCT medium (Ames, Elkhar, IN) then flash frozen in liquid nitrogen. All leprosy patients were recruited with approval from the Institutional Review Board of University of Southern California school of Medicine or the institutional ethics committee of Oswald Cruz Foundation, as well as the University of California, Los Angeles.

### Dual RNA sequencing of leprosy skin lesions

Frozen tissue sections of nine L-lep, nine RR, and six T-lep frozen skin lesions were cut into multiple ten micron sections and lysed in Qiagen RLT buffer with 0.1% β-mercaptoethanol (Invitrogen). To ensure efficient lysis of mycobacterial cell wall, lysates were homogenized with 0.1mm silica beads (MP Biomedicals, Santa Ana, CA) in the FastPrep3000 twice at a setting of 6.5 for 45s. RNA was extracted via Qiagen AllPrep Kit (Qiagen, Hilden, Germany). Ribosomal RNA was depleted with the Illumina Ribo-Zero Gold rRNA Removal Kit (Epidemiology) before sequencing library preparation with the Illumina Truseq Stranded Total RNA Sample Preparation Kit. Libraries were sequenced on the HiSeq2000 Sequencer (Illumina) at singleend 50 bp reads. Reads with poor quality were trimmed by T rim Galore! (http://www.bioinformatics.babraham.ac.uk/projects/trim_galore) and high quality reads were mapped to the hg19 human genome via STAR (Dobin et al., 2013). Unmapped reads after human genome alignment were mapped to the Br4923 *M. leprae* genome (Assembly ASM2668v1). The sum of exonic reads per gene were counted using HTSeq (Anders) using the GENCODE GRCh37-mapped version Release 24 (Harrow et al., 2012). Although, *M. leprae* 16S ribosomal RNA was depleted, a 96bp region (1343455-1343551) of the 23S rRNA remained abundant (Figure S2). Presumably, the approach to deplete rRNA did not include oligonucleotides that bound to these sequences and rRNA was fragmented. To confirm, the entire *M. leprae* 23S rRNA sequence and undepleted region were entered into the manufacturer website Epicentre Matchmaker (http://www.epibio.com/rnamatchmaker) and indicated “No Match” for binding of Ribozero depletion probes, while the well-depleted 16S rRNA sequence indicated a 65-85% match. Expression of *M. leprae* 23S rRNA and esxA were measured by quantitative real-time PCR KAPA SYBR FAST (MilliporeSigma, St Louis, MO) of cDNA synthesized from the same lesional RNA used in sequencing but before any ribosomal depletion. The relative quantities of the gene tested per sample were calculated against the GAPDH mRNA using the CT formula as previously described (Teles et al., 2013).

### Data analysis

Filtered read counts (containing genes with ≥ 5 reads in at least one patient group) from bacterial and human transcriptomes were normalized via DESeq2 (Love et al., 2014) using default parameters, except for basing scaling factors on only human genes. DESeq2 normalized gene counts and rlog-transformed counts were output and used for subsequent data analysis. Unsupervised hierarchical clustering was performed on rlog gene counts filtered for adequate expression of ≥ 5 rlog of gene in at least 20% of samples and filtered for variance by having a CV ≥ 0.2 rlog. Transcript abundance per gene was calculated by dividing the number of normalized read counts per gene by the gene length (exons only). *M. leprae* RNA abundance was calculated by taking the sum of the transcript abundance of all bacterial genes was divided by the total abundance of bacterial or human genes per sample. *M. leprae* transcriptional activity was calculated by dividing of the sum of the transcript abundance of all bacterial mRNA genes by the transcription abundance of 23S (based on the 1343455-1343551 region). A log transformation (log2(x+1)) of the *M. leprae* transcriptional activity was used for all analyses.

### SaVanT cell signature calculation

Cell type signatures were calculated using the web-based Signature Visualization Tool (SaVanT; http://newpathways.mcdb.ucla.edu/savant) (Lopez et al., 2017). In brief, SaVanT calculates a cell type score based on averaging a set of pre-calculated signature genes for each cell type which is proportional to the cell abundance within the sample. Human data from skin lesions (rlog-transformed gene counts) were input into SaVanT with the setting to use the top 50 genes per signature for score calculation. Since the output scores are in arbitrary expression units, the z-score was calculated per sample relative to all samples to depict the same scale for all signatures.

### Mycobacterial axenic culture experiments

*M. leprae* was grown in the footpad of nu/nu mice, as described previously (Lahiri et al., 2005) and harvested at log phase of growth and ethanol fixed *in situ* or cultured in NHDP medium for 48h or 96h post-harvest. Total RNA was extracted and rRNA depleted with Illumina Ribo-Zero Gold rRNA Removal Kit (Epidemiology). The Solid 5500 system was used for sequencing and quality control, alignment was performed by Maverix Biomics (San Mateo, CA). Gene counts (including 23S regional calculation) and transcriptional activity were calculated as described for lesional RNAseq analysis above. *M. tuberculosis* experimental data was derived from a re-analysis of previously published studies (Arnvig et al., 2011). Nutrient deprivation experiment was performed by growing *M. tuberculosis* H37Rv in Middlebrook 7H9 medium supplemented with 0.4% glycerol, 0.085% NaCl, 0.5% BSA, and 0.05% Tyloxapol. For nutrient depletion condition, exponentially growing bacteria were washed, resuspended in PBS supplemented with 0.025% Tyloxapol, and cultured for a further 24h. RNA was extracted and Illumina Sequencing libraries were constructed and sequenced as described (Cortes et al., 2013). In the exponential and stationary phase *M. tuberculosis* comparison, H37Rv was grown in Middlebrook 7H9 supplemented with 0.2% glycerol and 10%ADC. Exponential phase cultures were harvested at OD_600_ 0.6 to 0.8; stationary phase cultures were harvested one week after OD_600_ had reached 1.0. RNA was from three independent exponential phase cultures of *M. tuberculosis* and two stationary phase cultures were used to generate cDNA preparations that were then analyzed by Illumina-based sequencing as described (Arnvig et al., 2011).

### M. leprae infection of macrophages in vitro

Monocyte derived macrophages were derived from healthy human monocytes, purified by CD14+ microbead enrichment (Miltenyi Biotec, Bergisch Gladbach, Germany). Monocytes were differentiated into macrophages for 5 days in M-CSF 50ug/ml as previously described (Realegeno et al., 2016). Macrophages were infected at a multiplicity of infection of ten bacteria per cell from live *M. leprae* that was grown in the footpad of nu/nu mice, as described above. RNA was extracted by Qiagen AllPrep Kit (Qiagen, Hilden, Germany) at indicated timepoints. Ribosomal RNA was depleted, Illumina stranded libraries prepared, sequenced, and raw data analyzed in same method as leprosy lesions described above. All healthy human donors were recruited with approval from the Institutional Review Board of the University of California at Los Angeles.

### Weighted gene correlation analysis (WGCNA) and gene enrichment analysis

Module analysis of bacterial or human correlation networks was performed using Weighted Gene Correlation Analysis (WGCNA) (Langfelder and Horvath, 2008a). The fraction abundance of mRNA for *M. leprae* transcriptome was calculated by dividing the transcriptional abundance of each gene (DESeq2 normal counts/gene length) by the sum of all abundances of mRNA genes. Input for the *M. leprae* WGCNA was the mRNA fractional abundance with log-transformation (log2(x + 1)) and human WGCNA was rlog-transformed output from DESeq2 normalization. A signed gene correlation network was constructed using the “blockwiseModules()” function with a soft thresholding power of 12 for human and 16 for *M. leprae* and using a minimum module size of 50 genes, merged cut threshold of 0.3, and deepSplit of 0 for both analyses. The module eigengene, which represents a linear combination of genes that capture a large fraction of variance in each module, was calculated for each sample and correlated to *M. leprae* abundance or transcriptional activity per sample.

Bacterial module gene enrichment analysis was performed through hypergeometric overlap analysis of the Mycobrowser functional categories (Kapopoulou et al., 2011) for each gene. Further enrichment analysis of the *MLEPblue* module virulence genes was performed by the Database for Annotation, Visualization and Integrated Discovery (DAVID). Human module gene enrichment was performed using Gene Set Enrichment Analysis (GSEA) (Subramanian et al., 2005) of the Molecular Signatures Database (MSigDB) (Liberzon et al., 2011). Further enrichment analysis of the *HUMANgreenyellow* module was performed by Ingenuity Pathways Analysis (Qiagen, Hilden, Germany). Canonical pathway ‘Communication between Innate and Adaptive Immune Cells’ was edited to include only the portion of the pathway which had enriched genes and for visual clarity by the Pathway Designer function.

### Correlation and statistical analysis

Pearson correlation plots (Figure 4D and Figure 5A) were constructed using the ‘corrplot’ R package (R Core Team, 2013; Wei, 2017). Correlation significance test for association between paired samples was performed by the R function “cor.test()” using Pearson’s product moment correlation coefficient. The two-tailed student’s t-test was used when individual comparisons between two groups were performed. Individual details of statistical analyses are explained in the figure legends.

## References

Anders, S. (2010). HTSeq: Analysing high-throughput sequencing data with Python. URL http://www-huberemblde/users/anders/HTSeq/doc/overviewhtml.

Arnvig, K.B., Comas, I., Thomson, N.R., Houghton, J., Boshoff, H.I., Croucher, N.J., Rose, G., Perkins, T.T., Parkhill, J., Dougan, G. et al. (2011). Sequence-based analysis uncovers an abundance of non-coding RNA in the total transcriptome of Mycobacterium tuberculosis. PLoS Pathog 7, e1002342.

Asea, A., Rehli, M., Kabingu, E., Boch, J.A., Bare, O., Auron, P.E., Stevenson, M.A., and Calderwood, S.K. (2002). Novel signal transduction pathway utilized by extracellular HSP70: role of toll-like receptor (TLR) 2 and TLR4. J Biol Chem 277, 15028–15034.

Barton, R.P. (1975). Importance of nasal lesions in early lepromatous leprosy. Ann R Coll Surg Engl 57, 309–312.

Batoni, G., Maisetta, G., Florio, W., Freer, G., Campa, M., and Senesi, S. (1998). Analysis of the Mycobacterium bovis hsp60 promoter activity in recombinant Mycobacterium avium. FEMS Microbiol Lett 169, 117–124.

Bleharski, J.R., Li, H.Y., Meinken, C., Graeber, T.G., Ochoa, M.T., Yamamura, M., Burdick, A., Sarno, E.N., Wagner, M., Rollinghoff, M. et al. (2003). Use of genetic profiling in leprosy to discriminate clinical forms of the disease. Science 301, 1527–1530.

Boxx, G.M., and Cheng, G. (2016). The Roles of Type | Interferon in Bacterial Infection. Cell Host Microbe 19, 760–769.

Bulut, Y., Michelsen, K.S., Hayrapetian, L., Naiki, Y., Spallek, R., Singh, M., and Arditi, M. (2005). Mycobacterium tuberculosis heat shock proteins use diverse Toll-like receptor pathways to activate pro-inflammatory signals. J Biol Chem 280, 20961–20967.

Cangelosi, G.A., and Brabant, W.H. (1997). Depletion of pre-16S rRNA in starved Escherichia coli cells. J Bacteriol 179, 4457–4463.

Chuang, Y.M., Pinn, M.L., Karakousis, P.C., and Hung, C.F. (2018). Intranasal Immunization with DnaK Protein Induces Protective Mucosal Immunity against Tuberculosis in CD4-Depleted Mice. Frontiers in cellular and infection microbiology 8, 31.

Cortes, T., Schubert, O.T., Rose, G., Arnvig, K.B., Comas, I., Aebersold, R., and Young, D.B. (2013). Genome-wide mapping of transcriptional start sites defines an extensive leaderless transcriptome in Mycobacterium tuberculosis. Cell reports 5, 1121–1131.

Damron, F.H., Oglesby-Sherrouse, A.G., Wilks, A., and Barbier, M. (2016). Dual-seq transcriptomics reveals the battle for iron during Pseudomonas aeruginosa acute murine pneumonia. Proc Natl Acad Sci U S A 6, 39172.

Davis, G.L., Ray, N.A., Lahiri, R., Gillis, T.P., Krahenbuhl, J.L., Williams, D.L., and Adams, L.B. (2013). Molecular assays for determining Mycobacterium leprae viability in tissues of experimentally infected mice. PLoS Negl Trop Dis 7, e2404.

de Larranaga, G.F., Forastiero, R.R., Martinuzzo, M.E., Carreras, L.O., Tsariktsian, G., Sturno, M.M., and Alonso, B.S. (2000). High prevalence of antiphospholipid antibodies in leprosy: evaluation of antigen reactivity. Lupus 9, 594–600.

Dobin, A., Davis, C.A., Schlesinger, F., Drenkow, J., Zaleski, C., Jha, S., Batut, P., Chaisson, M., and Gingeras, T.R. (2013). STAR: ultrafast universal RNA-seq aligner. Bioinformatics 29, 15–21.

Dorner, M., Brandt, S., Tinguely, M., Zucol, F., Bourquin, J.P., Zauner, L., Berger, C., Bernasconi, M., Speck, R.F., and Nadal, D. (2009). Plasma cell toll-like receptor (TLR) expression differs from that of B cells, and plasma cell TLR triggering enhances immunoglobulin production. Immunology 128, 573–579.

Eng, R.H., Padberg, F.T., Smith, S.M., Tan, E.N., and Cherubin, C.E. (1991). Bactericidal effects of antibiotics on slowly growing and nongrowing bacteria. Antimicrob Agents Chemother 35, 1824–1828.

Fabri, M., Stenger, S., Shin, D.M., Yuk, J.M., Liu, P.T., Realegeno, S., Lee, H.M., Krutzik, S.R., Schenk, M., Sieling, P.A., et al. (2011). Vitamin D Is Required for IFN-gamma-Mediated Antimicrobial Activity of Human Macrophages. Sci Transl Med 3, 104ra102.

Giri, P.K., Kruh, N.A., Dobos, K.M., and Schorey, J.S. (2010). Proteomic analysis identifies highly antigenic proteins in exosomes from M. tuberculosis-infected and culture filtrate protein-treated macrophages. Proteomics 10, 3190–3202.

Gross, J.A., Johnston, J., Mudri, S., Enselman, R., Dillon, S.R., Madden, K., Xu, W., Parrish-Novak, J., Foster, D., Lofton-Day, C., et al. (2000). TACI and BCMA are receptors for a TNF homologue implicated in B-cell autoimmune disease. Nature 404, 995–999.

Harrow, J., Frankish, A., Gonzalez, J.M., Tapanari, E., Diekhans, M., Kokocinski, F., Aken, B.L., Barrell, D., Zadissa, A., Searle, S., et al. (2012). GENCODE: the reference human genome annotation for The ENCODE Project. Genome Res 22, 1760–1774.

Huang, d.W., Sherman, B.T., and Lempicki, R.A. (2009). Bioinformatics enrichment tools: paths toward the comprehensive functional analysis of large gene lists. Nucleic Acids Res 37, 1–13.

Iyer, A.M., Mohanty, K.K., van, E.D., Katoch, K., Faber, W.R., Das, P.K., and Sengupta, U. (2007). Leprosy-specific B-cells within cellular infiltrates in active leprosy lesions. Hum Pathol 38, 1065–1073.

Kapopoulou, A., Lew, J.M., and Cole, S.T. (2011). The MycoBrowser portal: a comprehensive and manually annotated resource for mycobacterial genomes. Tuberculosis (Edinburgh, Scotland) 91, 8—13.

Kester, J.C., and Fortune, S.M. (2014). Persisters and beyond: mechanisms of phenotypic drug resistance and drug tolerance in bacteria. Crit Rev Biochem Mol Biol 49, 91–101.

Kim, J., Gross, J.A., Dillon, S.R., Min, J.K., and Elkon, K.B. (2011). Increased BCMA expression in lupus marks activated B cells, and BCMA receptor engagement enhances the response to TLR9 stimulation. Autoimmunity 44, 69–81.

Kirchheimer, W.F., and Storrs, E.E. (1971). Attempts to establish the armadillo (Dasypus novemcinctus Linn.) as a model for the study of leprosy. I. Report of lepromatoid leprosy in an experimentally infected armadillo. Int J Lepr 39, 693–702.

Kralik, P., Nocker, A., and Pavlik, I. (2010). Mycobacterium avium subsp. paratuberculosis viability determination using F57 quantitative PCR in combination with propidium monoazide treatment. Int J Food Microbiol 141 Suppl 1, S80–S86.

Lahiri, R., Randhawa, B., and Krahenbuhl, J. (2005). Application of a viability-staining method for Mycobacterium leprae derived from the athymic (nu/nu) mouse foot pad. J Med Microbiol 54, 235–242.

Langfelder, P., and Horvath, S. (2008a). WGCNA: an R package for weighted correlation network analysis. BMC Bioinformatics 9, 559.

Langfelder, P., and Horvath, S. (2008b). WGCNA: an R package for weighted correlation network analysis. BMCBioinformatics 9, 559.

Liberzon, A., Subramanian, A., Pinchback, R., Thorvaldsdottir, H., Tamayo, P., and Mesirov, J.P. (2011). Molecular signatures database (MSigDB) 3.0. Bioinformatics 27, 1739–1740.

Lima, K.M., dos Santos, S.A., Santos, R.R., Brandao, I.T., Rodrigues, J.M., Jr., and Silva, C.L. (2003). Efficacy of DNA-hsp65 vaccination for tuberculosis varies with method of DNA introduction in vivo. Vaccine 22, 49–56.

Liu, P.T., Wheelwright, M., Teles, R., Komisopoulou, E., Edfeldt, K., Ferguson, B., Mehta, M.D., Vazirnia, A., Rea, T.H., Sarno, E.N., et al. (2012). MicroRNA-21 targets the vitamin D-dependent antimicrobial pathway in leprosy. Nat Med 18, 267–273.

Loizou, S., Singh, S., Wypkema, E., and Asherson, R.A. (2003). Anticardiolipin, anti-beta(2)-glycoprotein | and antiprothrombin antibodies in black South African patients with infectious disease. Ann Rheum Dis 62, 1106–1111.

Lopez, D., Montoya, D., Ambrose, M., Lam, L., Briscoe, L., Adams, C., Modlin, R.L., and Pellegrini, M. (2017). SaVanT: a web-based tool for the sample-level visualization of molecular signatures in gene expression profiles. BMC Genomics 18, 824.

Lorenzi, J.C., Trombone, A.P., Rocha, C.D., Almeida, L.P., Lousada, R.L., Malardo, T., Fontoura, I.C., Rossetti, R.A., Gembre, A.F., Silva, A.M., et al. (2010). Intranasal vaccination with messenger RNA as a new approach in gene therapy: use against tuberculosis. BMC Biotechnol 10, 77.

Love, M.I., Huber, W., and Anders, S. (2014). Moderated estimation of fold change and dispersion for RNA-seq data with DESeq2. Genome Biol 15, 550.

Lowrie, D.B., Tascon, R.E., Bonato, V.L., Lima, V.M., Faccioli, L.H., Stavropoulos, E., Colston, M.J., Hewinson, R.G., Moelling, K., and Silva, C.L. (1999). Therapy of tuberculosis in mice by DNA vaccination. Nature 400, 269–271.

Lu, L.L., Chung, A.W., Rosebrock, T.R., Ghebremichael, M., Yu, W.H., Grace, P.S., Schoen, M.K., Tafesse, F., Martin, C., Leung, V., et al. (2016). A Functional Role for Antibodies in Tuberculosis. Cell 167, 433–443.e414.

Lupoli, T.J., Fay, A., Adura, C., Glickman, M.S., and Nathan, C.F. (2016). Reconstitution of a Mycobacterium tuberculosis proteostasis network highlights essential cofactor interactions with chaperone DnaK. Proc Natl Acad Sci U S A 113, E7947–E7956.

Mackay, F., Woodcock, S.A., Lawton, P., Ambrose, C., Baetscher, M., Schneider, P., Tschopp, J., and Browning, J.L. (1999). Mice transgenic for BAFF develop lymphocytic disorders along with autoimmune manifestations. J Exp Med 190, 1697–1710.

Madigan, C.A., Cambier, C.J., Kelly-Scumpia, K.M., Scumpia, P.O., Cheng, T.Y., Zailaa, J., Bloom, B.R., Moody, D.B., Smale, S.T., Sagasti, A., et al. (2017a). A Macrophage Response to Mycobacterium leprae Phenolic Glycolipid Initiates Nerve Damage in Leprosy. Cell 170, 973–985.e910.

Madigan, C.A., Cameron, J., and Ramakrishnan, L. (2017b). A Zebrafish Model of Mycobacterium leprae Granulomatous Infection. J Infect Dis 216, 776–779.

Marsters, S.A., Yan, M., Pitti, R.M., Haas, P.E., Dixit, V.M., and Ashkenazi, A. (2000). Interaction of the TNF homologues BLyS and APRIL with the TNF receptor homologues BCMA and TACI. Curr Biol 10, 785–788.

Martin, C.J., Booty, M.G., Rosebrock, T.R., Nunes-Alves, C., Desjardins, D.M., Keren, I., Fortune, S.M., Remold, H.G., and Behar, S.M. (2012). Efferocytosis is an innate antibacterial mechanism. Cell Host Microbe 12, 289–300.

Martinez, A.N., Lahiri, R., Pittman, T.L., Scollard, D., Truman, R., Moraes, M.O., and Williams, D.L. (2009). Molecular determination of Mycobacterium leprae viability by use of real-time PCR. J Clin Microbiol 47, 2124–2130.

Modlin, R.L., Hofman, F.M., Taylor, C.R., and Rea, T.H. (1982). In situ characterization of T lymphocyte subsets in leprosy granulomas [letter]. Int J Lepr 50, 361–362.

Montoya, D., Cruz, D., Teles, R.M., Lee, D.J., Ochoa, M.T., Krutzik, S.R., Chun, R., Schenk, M., Zhang, X., Ferguson, B.G., et al. (2009). Divergence of macrophage phagocytic and antimicrobial programs in leprosy. Cell Host Microbe 6, 343–353.

Montoya, D., Inkeles, M.S., Liu, P.T., Realegeno, S., Teles, R.M.B., Vaidya, P., Munoz, M.A., Schenk, M., Swindell, W.R., Chun, R., et al. (2014). IL-32 is a molecular marker of a host defense network in human tuberculosis. Sci Transl Med 6, 250ra114.

Niemiec, M.J., Grumaz, C., Ermert, D., Desel, C., Shankar, M., Lopes, J.P., Mills, I.G., Stevens, P., Sohn, K., and Urban, C.F. (2017). Dual transcriptome of the immediate neutrophil and Candida albicans interplay. BMC Genomics 18, 696.

Nuss, A.M., Beckstette, M., Pimenova, M., Schmuhl, C., Opitz, W., Pisano, F., Heroven, A.K., and Dersch, P. (2017). Tissue dual RNA-seq allows fast discovery of infection-specific functions and riboregulators shaping host-pathogen transcriptomes. 114, E791–e800.

Ochoa, M.T., Teles, R., Haas, B.E., Zaghi, D., Li, H., Sarno, E.N., Rea, T.H., Modlin, R.L., and Lee, D.J. (2010). A role for interleukin-5 in promoting increased immunoglobulin M at the site of disease in leprosy. Immunology 131, 405–414.

Pittman, K.J., Aliota, M.T., and Knoll, L.J. (2014). Dual transcriptional profiling of mice and Toxoplasma gondii during acute and chronic infection. BMC Genomics 15, 806.

R Core Team (2013). R: A language and environment for statistical computing. (Vienna, Austria: R Foundation for Statistical Computing).

Realegeno, S., Kelly-Scumpia, K.M., Dang, A.T., Lu, J., Teles, R., Liu, P.T., Schenk, M., Lee, E.Y., Schmidt, N.W., Wong, G.C., et al. (2016). S100A12 Is Part of the Antimicrobial Network against Mycobacterium leprae in Human Macrophages. PLoS Pathog 12, e1005705.

Ridley, D.S. (1957). The use biopsies in therapeutic trials in leprosy. Trans R Soc Trop Med Hyg 51, 152–156.

Ridley, D.S., and Jopling, W.H. (1966). Classification of leprosy according to immunity. A five-group system. Int J Lepr 34, 255–273.

Rustad, T.R., Minch, K.J., Brabant, W., Winkler, J.K., Reiss, D.J., Baliga, N.S., and Sherman, D.R. (2013). Global analysis of mRNA stability in Mycobacterium tuberculosis. Nucleic Acids Res 41, 509–517.

Sille, F.C., Martin, C., Jayaraman, P., Rothchild, A., Fortune, S., Besra, G.S., Behar, S.M., and Boes, M. (2011). Requirement for invariant chain in macrophages for Mycobacterium tuberculosis replication and CD1d antigen presentation. Infect Immun 79, 3053–3063.

Souza, P.R., Zarate-Blades, C.R., Hori, J.I., Ramos, S.G., Lima, D.S., Schneider, T., Rosada, R.S., Torre, L.G., Santana, M.H., Brandao, I.T. et al. (2008). Protective efficacy of different strategies employing Mycobacterium leprae heat-shock protein 65 against tuberculosis. Expert Opin Biol Ther 8, 1255–1264.

Stanley, S.A., Johndrow, J.E., Manzanillo, P., and Cox, J.S. (2007). The Type | IFN response to infection with Mycobacterium tuberculosis requires ESX-1-mediated secretion and contributes to pathogenesis. J Immunol 178, 3143–3152.

Subramanian, A., Kuehn, H., Gould, J., Tamayo, P., and Mesirov, J.P. (2007). GSEA-P: a desktop application for Gene Set Enrichment Analysis. Bioinformatics 23, 3251–3253.

Subramanian, A., Tamayo, P., Mootha, V.K., Mukherjee, S., Ebert, B.L., Gillette, M.A., Paulovich, A., Pomeroy, S.L., Golub, T.R., Lander, E.S., et al. (2005). Gene set enrichment analysis: a knowledge-based approach for interpreting genome-wide expression profiles. Proc Natl Acad Sci U S A 102, 15545–15550.

Tascon, R.E., Colston, M.J., Ragno, S., Stavropoulos, E., Gregory, D., and Lowrie, D.B. (1996). Vaccination against tuberculosis by DNA injection. Nature Med 2, 888–892.

Teles, R.M., Graeber, T.G., Krutzik, S.R., Montoya, D., Schenk, M., Lee, D.J., Komisopoulou, E., Kelly-Scumpia, K., Chun, R., Iyer, S.S., et al. (2013). Type | interferon suppresses type II interferon-triggered human anti-mycobacterial responses. Science 339, 1448–1453.

Thanert, R., Goldmann, O., Beineke, A., and Medina, E. (2017). Host-inherent variability influences the transcriptional response of Staphylococcus aureus during in vivo infection. Nature communications 8, 14268.

Vargas-Romero, F., Guitierrez-Najera, N., Mendoza-Hernandez, G., Ortega-Bernal, D., Hernandez-Pando, R., and Castanon-Arreola, M. (2016). Secretome profile analysis of hypervirulent Mycobacterium tuberculosis CPT31 reveals increased production of EsxB and proteins involved in adaptation to intracellular lifestyle. Pathogens and disease 74.

Wei, T.S., Villam (2017). R package “corrplot“: Visualization of a Correlation Matrix (Version 0.84).

Wesolowska-Andersen, A., Everman, J.L., Davidson, R., Rios, C., Herrin, R., Eng, C., Janssen, W.J., Liu, A.H., Oh, S.S., Kumar, R. et al. (2017). Dual RNA-seq reveals viral infections in asthmatic children without respiratory illness which are associated with changes in the airway transcriptome. Genome Biol 18, 12.

Westermann, A.J., Forstner, K.U., Amman, F., Barquist, L., Chao, Y., Schulte, L.N., Muller, L., Reinhardt, R., Stadler, P.F., and Vogel, J. (2016). Dual RNA-seq unveils noncoding RNA functions in host-pathogen interactions. Nature 529, 496–501.

Yamamura, M., Uyemura, K., Deans, R.J., Weinberg, K., Rea, T.H., Bloom, B.R., and Modlin, R.L. (1991). Defining protective responses to pathogens: cytokine profiles in leprosy lesions. Science 254, 277–279.

Young, D., Lathigra, R., Hendrix, R., Sweetser, D., and Young, R.A. (1988). Stress proteins are immune targets in leprosy and tuberculosis. Proc Natl Acad Sci U S A 85, 4267–4270.

Zhang, J., Roschke, V., Baker, K.P., Wang, Z., Alarcon, G.S., Fessler, B.J., Bastian, H., Kimberly, R.P., and Zhou, T. (2001). Cutting edge: a role for B lymphocyte stimulator in systemic lupus erythematosus. J Immunol 166, 6–10.

Zimmermann, M., Kogadeeva, M., Gengenbacher, M., McEwen, G., Mollenkopf, H.J., Zamboni, N., Kaufmann, S.H.E., and Sauer, U. (2017). Integration of Metabolomics and Transcriptomics Reveals a Complex Diet of Mycobacterium tuberculosis during Early Macrophage Infection. mSystems 2.

Zimmermann, N., Thormann, V., Hu, B., Kohler, A.B., Imai-Matsushima, A., Locht, C., Arnett, E., Schlesinger, L.S., Zoller, T., Schurmann, M., et al. (2016). Human isotype-dependent inhibitory antibody responses against Mycobacterium tuberculosis. EMBO Mol Med 8, 1325–1339.

